# Impact of Amazonian Dance on Speech Performance in People with Parkinson’s Disease

**DOI:** 10.64898/2026.01.12.699086

**Authors:** Raquel Arigony Prates, Carlos Cristiano Espedito Guzzo, Maria Vitória Andreazza Duarte, Alberto Muñoz Sánchez, Patrícia Pauli, Flávia Gomes Martinez, Elren Passos-Monteiro, Adolfo M. García, Aline Nogueira Haas, Constantina Theofanopoulou

## Abstract

Dance-based interventions have consistently been shown to improve limb motor function in Parkinson’s disease (PD), yet their potential impact on other motor domains, particularly those supporting laryngeal-orofacial control needed for speech production, remains largely unexplored. Beyond motor speech functions, dance may also influence higher-order language processes, including semantic organization. To test these hypotheses, we conducted a 12-week randomized trial comparing an Amazonian Dance intervention to a matched-physical intensity control condition (Nordic Walking), incorporating automated speech and language analysis to provide objective, fine-grained quantification of communication outcomes. Participants in the dance arm showed significant improvements in prosody (Main Tone), voice quality (Harmonic to Noise Ratio), and semantic organization (Granularity), whereas the walking group showed declines in these metrics. The dance-related gains in prosody and semantics remained significant even after adjusting for demographic, cognitive, and clinical covariates. These findings suggest that dance may enhance both speech-motor and higher-order language functions in PD, potentially through mechanisms such as auditory-motor coupling, improved internal timing, and the engagement of overlapping neural substrates between dance and speech and/or language.

## Introduction

Parkinson’s disease (PD) is one of the fastest growing neurodegenerative disorders, currently affecting over 9 million people worldwide^1–3^, while its prevalence continues to rise with population aging^4^. In Brazil, a population-based study estimated a prevalence of 3.3% of PD among people older than 60^5^. People with Parkinson’s (PwP) typically display cardinal motor impairments (e.g., resting tremor, postural instability, and bradykinesia) alongside diverse cognitive and non-motor symptoms (e.g., cognitive decline, mood disturbance, and sleep disorders)^3,6–7^. These interacting deficits collectively lead to progressive functional decline and loss of autonomy that negatively impact quality of life and well-being.

Despite substantial research on gait and limb motor function, speech remains one of the least addressed motor symptoms in PD, even though it affects up to 89% of cases^8–9^. These deficits range from strictly motor-speech production deficits, such as reduced pitch variation, monotonous prosody, and increased pausing, to broader linguistic (non-motor) disturbances, including impoverished lexical retrieval, impaired action-verb generation, and simplified syntax^10–12^. These deficits compromise intelligibility and self-expression, contributing to social withdrawal and decreased communicative confidence. Considering its reliance on finely tuned motor coordination, speech production has been proposed as a sensitive biomarker of disease progression^13–16^ with recent machine-learning approaches demonstrating high diagnostic accuracy based on acoustic and linguistic features^17^.

While various forms of speech and language therapy (SLT), such as the Lee Silverman Voice Treatment (LSVT), have been widely used to address PD-related speech deficits, the robustness of evidence supporting their long-term efficacy remains limited^18^. Moreover, PwP have frequently expressed dissatisfaction with SLT methods that rely heavily on repetitive drills and isolated motor exercises, emphasizing instead the need for approaches centered on functional and communicative expressiveness^19–20^. Another limitation lies in the demanding intensity of many SLT protocols, such as LSVT-based regimens requiring four sessions per week^21^, which may not be feasible for many older or mobility-limited individuals. These limitations highlight the necessity of developing innovative, motivating, and ecologically valid interventions.

Dance-based interventions have emerged as promising non-pharmacological strategies for PwP, integrating rhythmic movement, social engagement and emotional expression that may benefit both motor and non-motor domains^22–24^. Numerous studies have consistently demonstrated improvements in balance, gait, and overall motor coordination following dance training^23–33^. Beyond limb locomotion, dance has been proposed to engage motor brain circuits that control the laryngeal and articulatory muscles involved in speech production, such as the primary motor cortex, as well as frontal and basal ganglia circuits implicated in higher-order aspects of language production (e.g., word retrieval)^34^. In line with this hypothesis, prior studies have reported enhanced verbal fluency, word recall, and word recognition following dance-based interventions, among other improvements^35–39^. However, the assessment tools used in these studies have generally lacked the granularity to disentangle whether these improvements primarily reflect changes in speech-motor control or in non-motor, language-specific processes—for instance, whether reduced word production arises from impaired memory retrieval or from difficulty in articulation—and have often relied on the subjectivity of human raters.

Building on this emerging evidence, we sought to bridge these gaps by examining whether a dance-based intervention (Amazonian Dance), compared with Nordic Walking, could produce measurable changes in speech-motor and/or language (non-motor) dimensions in PwP, using objective Automated Speech and Language Analysis (ASLA) features. Specifically, we employed the Toolkit to Examine Lifelike Language (TELL) app, which provides high-resolution acoustic and semantic metrics capable of parsing speech parameters that predominantly reflect motor control—such as tone, speech rate, average pause duration, and shimmer—from those indexing non-motor, language-specific processes, such as lexical-semantic precision and clustering in verbal fluency^40–41^. Further, echoing studies demonstrating the value of culturally adapted interventions in enhancing therapeutic engagement and efficacy^42^, the present study implemented the Amazonian Dance protocol, a culturally grounded approach derived from Northern Brazilian traditions, combining rhythm, storytelling, and symbolic movement^43^. Such culturally resonant frameworks may address the unique needs and experiences of PwP from diverse cultural backgrounds, promoting equity, reducing healthcare disparities, and ensuring culturally sensitive and patient-centered approaches that optimize quality of life. With this design, we hypothesized that participation in the Amazonian Dance intervention would lead to significant improvements in selected speech and language metrics, relative to the control condition.

## Results

The primary aim of this randomized controlled study was to determine whether a 12-week Amazonian Dance intervention, compared with an active control condition (Nordic Walking), produced measurable pre–post changes in speech-motor and/or language-related features in PwP. The Amazonian Dance intervention was implemented at two sites: a Dance South group (DS), comprising participants from Southern Brazil where identification with Amazonian dance traditions is less prevalent, and a Dance North group (DN), comprising participants from Northern Brazil who more strongly identify with these regional cultural practices. A total of 66 individuals were screened for eligibility, of whom 56 were enrolled and randomized into three groups (Dance South, Dance North, and Nordic Walking); attrition due to access difficulties, health issues, or incomplete post-intervention assessments resulted in a total of 36 participants with complete speech recordings: 20 participants in Dance South, 7 in Dance North, and 9 in the Nordic Walking group (**Figure 1**). The demographic and clinical characteristics of the analyzed sample demonstrated heterogeneity across groups in sex distribution, race/ethnicity, cognitive performance, disease severity, and years of education, underscoring the importance of subsequent covariate-adjusted analyses (**Table 1**).

**Figure 1.**
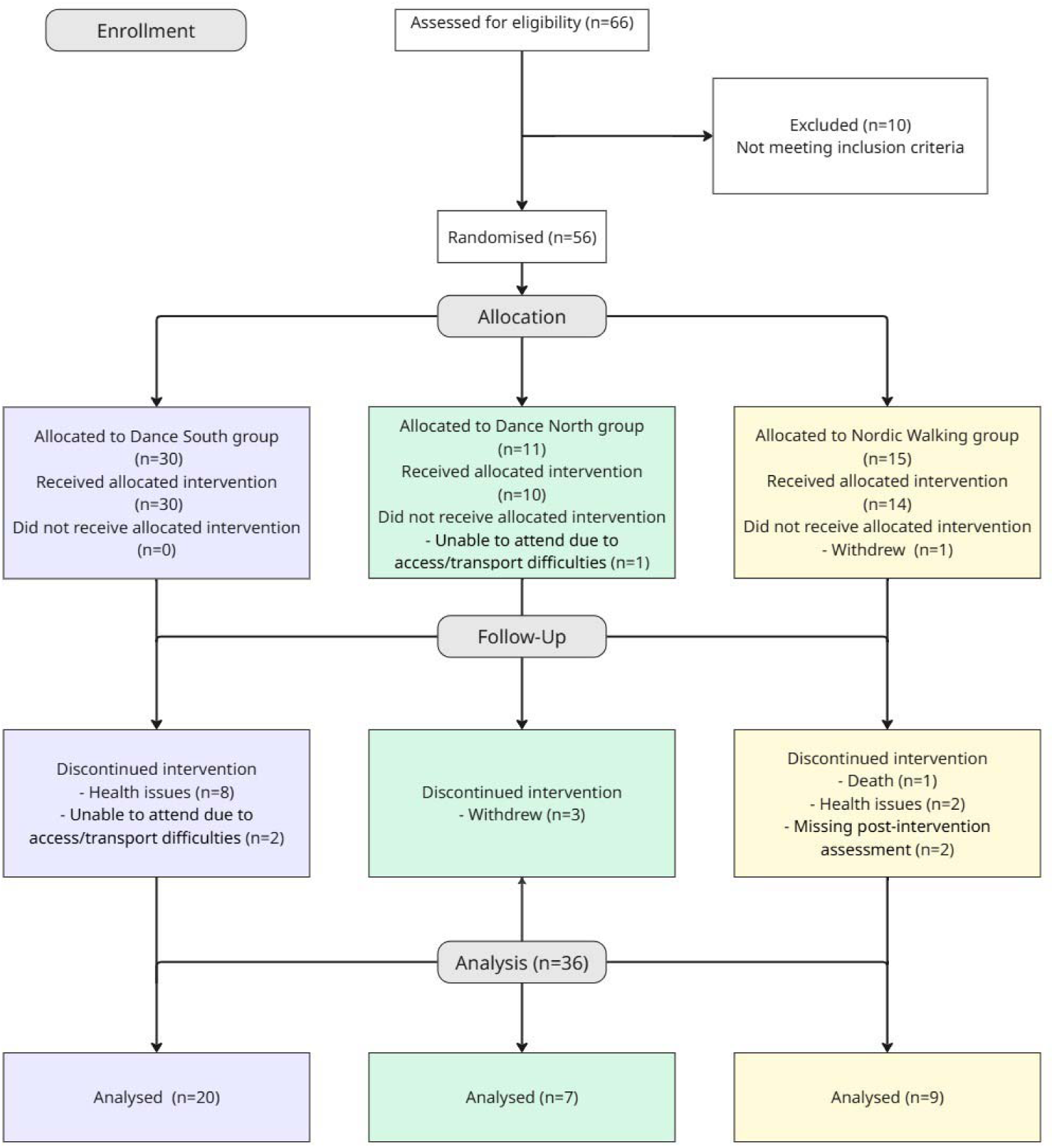
Participant flow diagram. Flowchart depicting screening, randomization, intervention adherence, follow-up, and inclusion in the final analysis. Of the 66 individuals assessed for eligibility, 56 met the inclusion criteria and were randomized to one of three groups: Dance South (n = 30), Dance North (n = 11), or Nordic Walking (n = 15). All participants who began the intervention were monitored throughout the 12-week program. Reasons for non-initiation or discontinuation included access difficulties, health issues, withdrawal, missing post-intervention assessments, and one death that occurred during the study period. A total of 36 participants (20 in Dance South, 7 in Dance North, and 9 in the Nordic Walking group) completed both pre- and post-intervention speech assessments and were included in the final analyses.

**Table 1.**
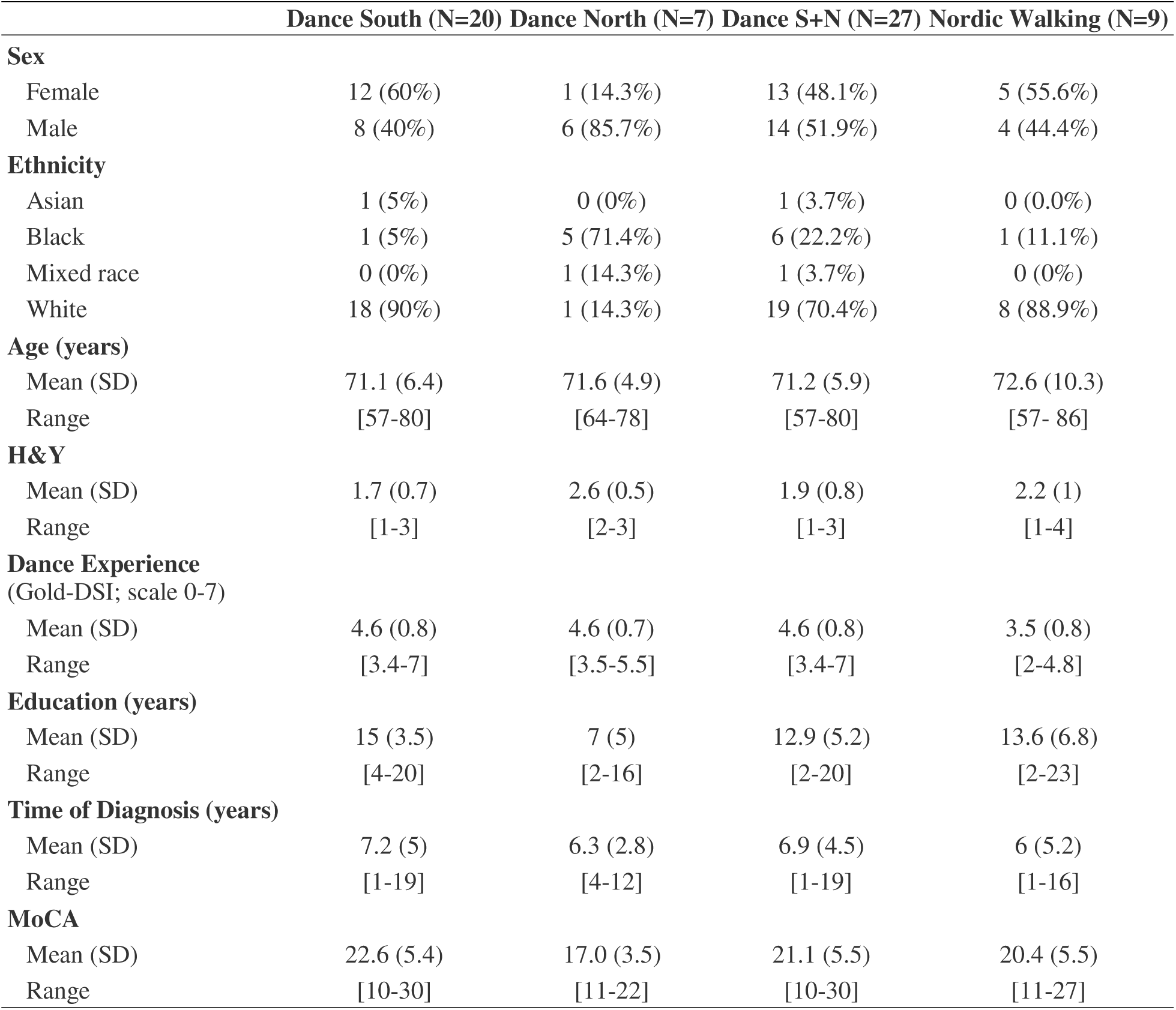
Participants’ demographic and clinical characteristics. Values represent the number and percentage of participants for categorical variables and mean (standard deviation) with range for continuous variables. The table summarizes baseline characteristics for participants who completed both pre- and post-intervention speech assessments in the Dance South (n = 20), Dance North (n = 7), Dance South +North (S+N) (n=27), and Nordic Walking (n = 9) groups. Ethnicity categories reflect self-identification using standard Brazilian census classifications. Dance experience was assessed using the Brazilian adaptation of the Goldsmiths Dance Sophistication Index (Gold-DSI; scale 0-7). MoCA scores were interpreted according to education-adjusted Brazilian cutoffs. H&Y = Hoehn and Yahr disease severity scale.

|  | Dance South (N=20) | Dance North (N=7) | Dance S+N (N=27) | Nordic Walking (N=9) |
| --- | --- | --- | --- | --- |
| **Sex** |  |  |  |  |
| Female | 12 (60%) | 1 (14.3%) | 13 (48.1%) | 5 (55.6%) |
| Male | 8 (40%) | 6 (85.7%) | 14 (51.9%) | 4 (44.4%) |
| **Ethnicity** |  |  |  |  |
| Asian | 1 (5%) | 0 (0%) | 1 (3.7%) | 0 (0.0%) |
| Black | 1 (5%) | 5 (71.4%) | 6 (22.2%) | 1 (11.1%) |
| Mixed race | 0 (0%) | 1 (14.3%) | 1 (3.7%) | 0 (0%) |
| White | 18 (90%) | 1 (14.3%) | 19 (70.4%) | 8 (88.9%) |
| **Age (years)** |  |  |  |  |
| Mean (SD) | 71.1 (6.4) | 71.6 (4.9) | 71.2 (5.9) | 72.6 (10.3) |
| Range | [57-80] | [64-78] | [57-80] | [57- 86] |
| **H&Y** |  |  |  |  |
| Mean (SD) | 1.7 (0.7) | 2.6 (0.5) | 1.9 (0.8) | 2.2 (1) |
| Range | [1-3] | [2-3] | [1-3] | [1-4] |
| **Dance Experience**<br>(Gold-DSI; scale 0-7) |  |  |  |  |
| Mean (SD) | 4.6 (0.8) | 4.6 (0.7) | 4.6 (0.8) | 3.5 (0.8) |
| Range | [3.4-7] | [3.5-5.5] | [3.4-7] | [2-4.8] |
| **Education (years)** |  |  |  |  |
| Mean (SD) | 15 (3.5) | 7 (5) | 12.9 (5.2) | 13.6 (6.8) |
| Range | [4-20] | [2-16] | [2-20] | [2-23] |
| **Time of Diagnosis (years)** |  |  |  |  |
| Mean (SD) | 7.2 (5) | 6.3 (2.8) | 6.9 (4.5) | 6 (5.2) |
| Range | [1-19] | [4-12] | [1-19] | [1-16] |
| **MoCA** |  |  |  |  |
| Mean (SD) | 22.6 (5.4) | 17.0 (3.5) | 21.1 (5.5) | 20.4 (5.5) |
| Range | [10-30] | [11-22] | [10-30] | [11-27] |

Speech features were extracted from two tasks, called “Animal Fluency” and “Routine Description”, administered at baseline and post-intervention. The Animal Fluency task required participants to name as many animals as possible within 60 seconds and probes semantic memory and lexical retrieval, whereas the Routine Description task elicited at least 60 seconds of spontaneous connected speech by asking participants to describe their typical morning routine. For each task, pre–post change scores (Δ = post − pre) were computed for all acoustic and semantic metrics. Analyses were conducted at multiple levels: within-group pre–post comparisons were first performed separately for each task and subsequently for composite scores averaged across the two tasks; between-group comparisons then tested differences in change values across all group contrasts, with false discovery rate (FDR) correction applied throughout. Finally, covariate models were used to assess whether observed group effects persisted after adjustment for demographic, cognitive, and clinical factors.

### Within-group analyses per task

We used within-group comparisons as an initial indication of how each intervention affected speech-motor and language features relative to participants’ own baselines (**Tables S1, S2**). In the Animal Fluency task, the Nordic Walking group showed three significant declines in both speech-motor and semantic features: Average Harmonics-to-Noise Ratio (HNR) decreased (Δ = −0.342, *p* = .0237), indicating reduced vocal harmonicity; Speech Rate decreased (Δ = −0.346, *p* = .0175), consistent with slower word production; and Granularity decreased (Δ = −0.596, *p* = .0488), reflecting less precise semantic content. In contrast, the Dance North group showed a significant increase in Granularity (Δ = +1.301, *p* = .0446), suggesting propensity towards more fine-grained conceptual processing within the Animal Fluency task. No other within-group changes reached significance in this task.

In the Routine task, the Dance North group exhibited a large and significant decrease in Main Tone (Δ = −18.337, *p* = .00470), a metric where lower values indicate beneficial shifts in predominant vocal pitch. The Dance North group also showed a significant change in Average pause duration (Δ = +0.269, *p* = .0156). Since in this feature shorter pauses signify improvement, this positive Δ reflects lengthened pauses (i.e., movement in the non-beneficial direction). No other within-group changes reached *p* < .05 in the Routine task. None of these within-group task-level effects survived FDR at q < .05, although Main Tone in the Dance North group (Routine) approached that threshold (q = .0564). Overall, these tests enabled us to detect task-specific changes in each group before evaluating between-group differences.

### Within-group analyses on composite scores

Composite analyses, which average each speech metric across both Animal Fluency and Routine, provide a task-independent estimate of overall pre−post change within each intervention arm (**Tables S3, S4**). With this approach, we aimed to reduce task-specific variability and detect broad intervention-related effects. The Nordic Walking group showed a significant composite decline in Average HNR (Δ = −0.294, *p* = .0134), consistent with worsening voice quality. The Dance North group showed significant composite improvement in Granularity (Δ = +0.661, *p* = .0494) and a large composite decrease in Main Tone (Δ = −12.455, *p* = .00775), again consistent with enhanced semantic output and beneficial prosodic change. A trend was observed for the Nordic Walking group Speech Rate (Δ = −0.187, *p* = .0509), suggesting a small decline in fluency that did not reach conventional significance. No composite within-group effects survived FDR at q < .05, although Main Tone (Dance North) approached that level (q ≈ .093). Taken together, the composite analyses indicate that the most consistent within-group changes were declines in vocal harmonicity for the Nordic Walking group and improvements in semantic precision and prosodic pitch for the Dance North group.

**Figure 2.**
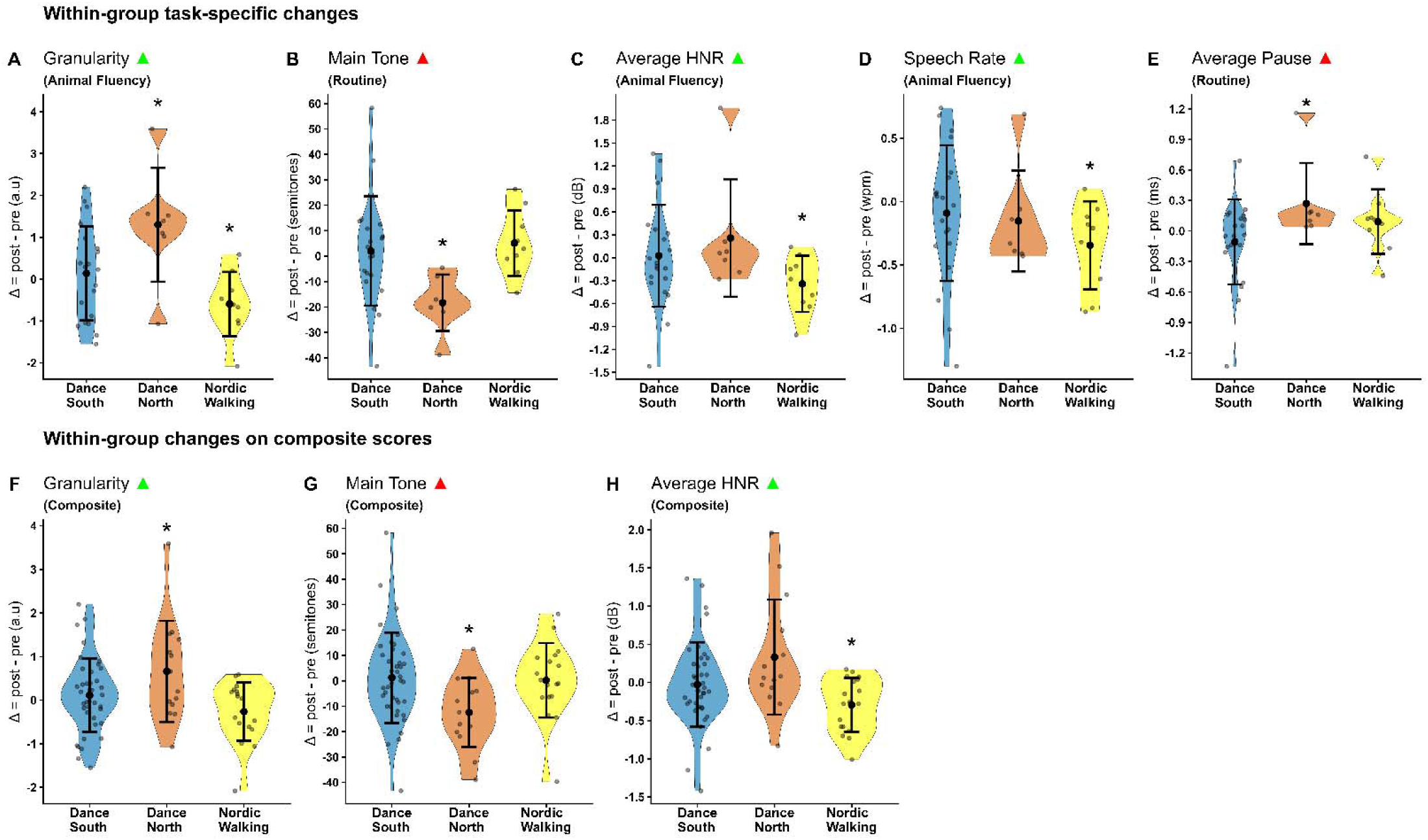
**Within-group changes (**Δ **= post–pre) in key speech and language metrics across the three intervention arms.** Violin plots display the distribution of individual-level change scores for each metric, separately for Dance South (blue), Dance North (orange), and Nordic Walking (yellow). Solid big points represent group means, the internal tick marks indicate the standard deviation, the widest part of the violin corresponds to the median, the upper and lower ends represent the first and third quartiles (Q1 and Q3), and jitter points show the raw sample size. Positive or negative Δ values reflect improvement or worsening depending on the metric’s directionality (e.g., higher Granularity and HNR indicate improvement; lower Average Pause Duration indicates improvement). Green arrows = the higher direction indicates improvement; red arrows = the higher direction indicates deterioration. Y-axis ranges differ across panels to reflect the natural scale of each metric (ms, semitones, etc). Granularity is plotted in arbitrary units (a.u.), Average Pause in milliseconds (ms); Speech Rate in words per minute (wpm), Average HNR in decibels (dB), and Main Tone in semitones. Asterisks denote statistically significant within-group pre–post changes: *p < .05 (uncorrected). n values per group: Dance South = 20; Dance North = 7; Nordic Walking = 9.

### Between-group analyses per task

We then ran between-group contrasts at the task level to directly compare how each intervention influenced specific speech features relative to the other arms (**Table S5**). In Animal Fluency, Granularity favored dance over Nordic Walking in two significant contrasts: Dance South + Dance North vs Nordic Walking (*p* = .00769, *g* ≈ 0.86; mean Δ dance = +0.439 vs Nordic walking = −0.596) and Nordic Walking vs Dance North (*p* = .00929, *g* ≈ −1.69; Nordic walking = −0.596 vs Dance North = +1.301), indicating better content organization in the dance arms. We also noted trend-level (*p* < .06) contrasts, with all changes being beneficial for the dance groups, in the following tasks: Average HNR (Dance South + Dance North vs Nordic Walking; *p* = .0506); Granularity (Dance South vs Nordic Walking; *p* = .0532), and Average HNR (Nordic Walking vs Dance North; *p* = .0549). All these between-group task-level effects did not survive FDR correction.

In the Routine task, Main Tone showed the most robust between-group differences favoring the Dance North group: Nordic Walking vs Dance North was significant and FDR-surviving (*p* = .00161, q = .0194, Hedges’ *g* = 1.83), with Nordic Walking showing a small positive Δ (+5.11) and Dance North a large negative change (−18.34), indicating improved prosody for Dance North. The contrast between Dance South and Dance North was also significant (*p* = .00448, q = .0537) for this feature, with Dance North showing improvement (Δ = −18.34) relative to no change in Dance South (Δ = +2.00). Further in this task, Average pause duration differed between Dance North and Dance South (*p* = .0379, rank-biserial *r* ≈ .543), with Dance South showing shorter pauses (Δ = −0.107) and Dance North longer pauses (Δ = +0.269). Overall, the task-level analyses indicate that the strongest between-group divergences occurred in prosodic pitch (Routine/Main Tone) and semantic detail (Animal Fluency/Granularity), with dance groups showing movement toward lower predominant pitch and higher granularity relative to Nordic Walking.

### Between-group analyses on composite scores

Similar to the within-group analyses, we then analyzed between-group comparisons in composite scores that allowed us to examine differences in overall post-pre change (Δ) across intervention arms by averaging each speech metric across both tasks, providing a broader estimate of intervention effects that is less influenced by task-specific variability. On composites, the clearest differences again favored dance arms, particularly Dance North, against Nordic Walking. Main Tone showed changes in the Dance South vs Dance North (*p* = .00886, *g* ≈ 0.93; Dance South Δ ≈ +1.21 vs Dance North Δ ≈ −12.46) and Nordic Walking vs Dance North comparisons (*p* = .0321, *g* ≈ 1.08; Nordic walking Δ ≈ +0.21 vs Dance North Δ ≈ −12.46), reflecting greater prosodic improvements in Dance North. Granularity was higher in dance vs Nordic walking (Nordic walking vs Dance North, *p* = .0143, *g* ≈ −1.50; Dance South + Dance North vs Nordic Walking, *p* = .0198, *g* ≈ 0.78), demonstrating more fine-grained semantic output for the dance arms. Average HNR (Dance South + Dance North vs Nordic walking) also favored the combined dance groups over Nordic walking (*p* = .0198; dance Δ ≈ +0.065 vs Nordic walking Δ ≈ −0.262), consistent with better average voice quality after dance. None of these composite between-group effects survived FDR at q < .05, but the pattern was internally consistent with the task-level results and with the within-group changes. Overall, the composite analyses reinforce that the largest between-group differences were observed in prosodic pitch, semantic detail, and vocal harmonicity, with the dance groups, particularly Dance North, showing more favorable change than Nordic Walking.

**Figure 3.**
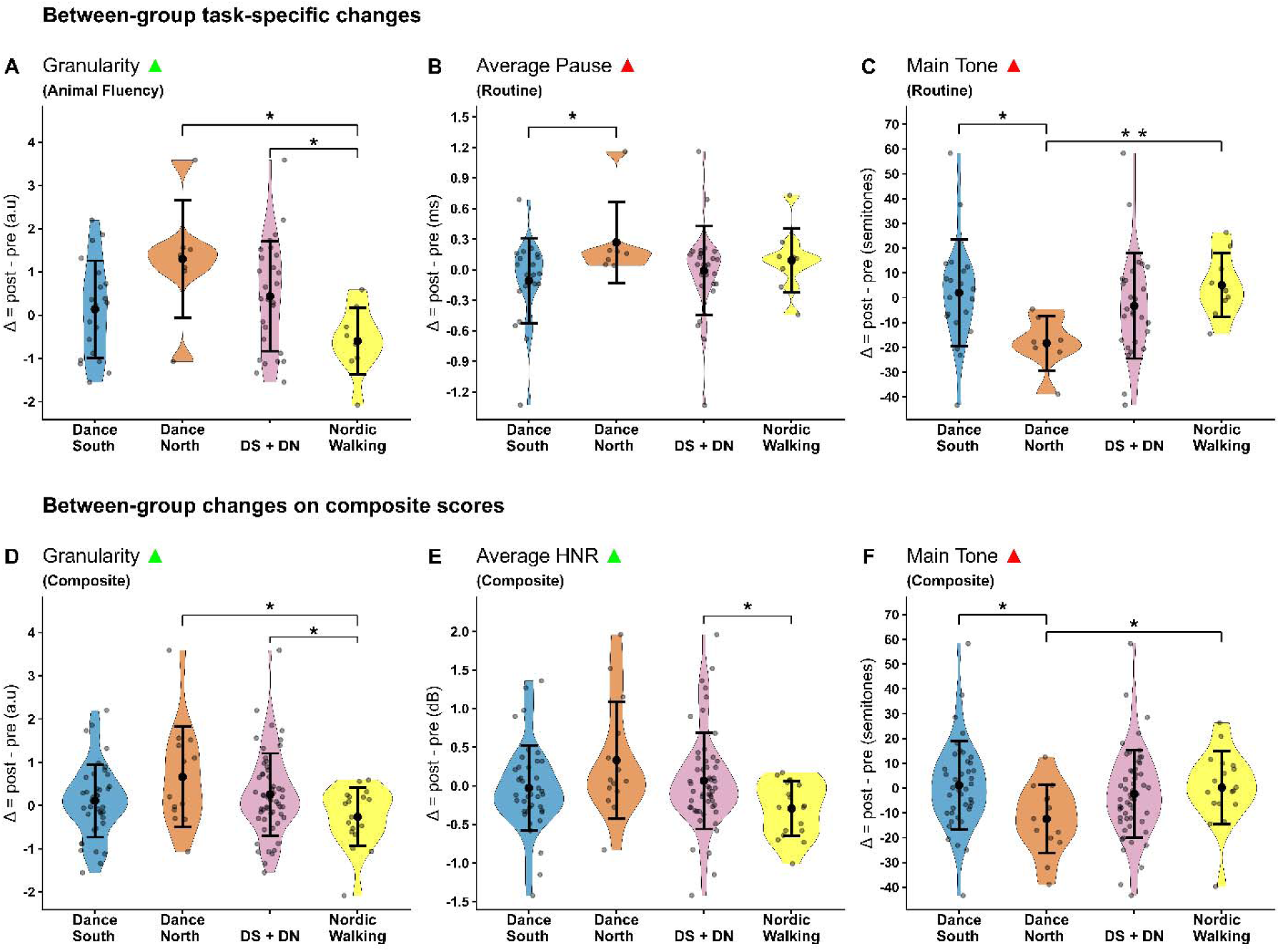
**Between-group differences in pre–post change (**Δ**) across speech and language metrics.** Violin plots depict the distribution of individual Δ values (post–pre) for each intervention arm: Dance South (blue), Dance North (orange), pooled Dance South and North (DS+DN; pink), and Nordic Walking (yellow). Each panel corresponds to a metric-task combination that showed significant between-group contrasts in the statistical analyses. Positive Δ values reflect increases and negative Δ reflect decreases in the feature; whether higher or lower values indicate improvement depends on the metric (e.g., increases in Granularity or HNR reflect improvement, whereas decreases in Main Tone reflect improvement). Green arrows = the higher direction indicates improvement; red arrows = the higher direction indicates deterioration. Big solid points denote group means, the internal tick marks indicate the standard deviation, the widest part of the violin corresponds to the median, the upper and lower ends represent the first and third quartiles (Q1 and Q3), and jitter points show the raw sample size. Brackets indicate significant between-group contrasts, where *p < .05 (uncorrected) and **q < .05 (FDR corrected). Y-axis ranges differ across panels to reflect the natural scale of each metric.

### Covariate analyses

To assess whether the observed group differences could be explained by demographic or clinical characteristics, we ran multivariable ANCOVA models including Group and seven covariates (Sex, Ethnicity, Dance experience, MoCA, Years of education, Age, and Hoehn and Yahr Scale) for every speech metric (**Tables S7-S10**). Even after adjustment, Group remained a significant predictor (p < .05) for a focused set of features, confirming that the intervention effects are not reducible to background characteristics. In Animal Fluency, Group effects survived for Average pause duration, Pause duration variability, Granularity, and Main tone. In Routine, Group remained significant for Ongoing Semantic Variability (OSV), Shimmer, Standard deviation HNR, and Main Tone. These models showed medium effect sizes (partial η² ≈ .10–.20), and several of them aligned directly with our primary intervention findings. Specifically, Main Tone (Routine), the clearest between-group separation in the task-level analysis, remained significant after covariates, and at the task level, Nordic Walking vs Dance North was also FDR-significant (q = .019). Granularity in the Animal Fluency task also emerged both as a Group-surviving effect in the adjusted models and as a robust between-group contrast (Groups Dance South + Dance North vs Nordic walking) in the unadjusted task-level analysis. By contrast, Average HNR (Dance South + Dance North vs Nordic Walking) in the composite between-group analysis, although favoring the dance arms before adjustment, did not survive as a Group effect once covariates were included, indicating that its variance was partially accounted for by demographic or clinical factors. Thus, only the effects that remained significant after covariate adjustment can be interpreted as intervention-related, whereas effects that did not survive adjustment are partly explained by underlying demographic or clinical variation.

### Influence and direction of specific covariates

Because the three intervention groups differed in demographic and clinical characteristics (**Table 1**), and because these characteristics may influence speech production in PD, we examined how each covariate related to patterns of speech change across metrics and tasks. This covariate-level analysis (**Table S7-S10**) clarifies whether observed improvements or declines might be attributable to specific background features rather than to the interventions themselves, and it helps identify subgroups that may differ in responsiveness.

Across demographic factors, ethnicity showed the broadest association footprint, with significant effects (p < .05) for 15 metrics (**Table S8**), especially timing/variability features such as Average pause duration and Pause duration variability in both tasks, and voice stability measures like Standard deviation HNR in Routine. Descriptively, Asian and White participants tended to show positive Δ across multiple metrics (Granularity, Speech rate, HNR, Pause variability), whereas Black and Brown/Mixed participants showed flatter or negative Δ overall, sometimes with strong gains on single metrics (Granularity, HNR) but declines on others (Speech rate, Pause variability). This suggests that the magnitude and consistency of speech gains varied by ethnic background, with more robust effects for some subgroups. Sex also influenced several metrics, with female participants showing larger positive Δ on content and prosodic features (Granularity, HNR, Standard deviation HNR), while males tended toward stable or negative Δ, suggesting that women contributed more strongly to the overall gains observed in the dance groups. Age showed significant associations with 9 metrics, with older participants demonstrating smaller improvements—or slight declines—in voice-quality measures such as HNR and increased variability, consistent with known age-related reductions in vocal stability that may blunt responsiveness to intervention.

Clinical and cognitive covariates also exhibited domain-specific patterns. H&Y stage was also widely associated (11 metrics), with participants at more advanced stages showing larger Δ on content/timing metrics such as Granularity and Pause duration variability, but smaller or negative Δ on prosody and voice stability features such as OSV and Shimmer. This indicates that disease severity modulated the profile of gains: while advanced-stage participants could still improve on content and timing, they showed less stable or even deteriorating performance on voice-quality measures. Dance experience predicted stronger gains in voice quality, especially higher HNR. MoCA showed smaller but interpretable effects, with higher MoCA scores linked to better speech (Syllable duration variability, Average syllable duration) and semantic (Granularity) scores, suggesting that stronger baseline cognition supported improvement in these features. Lastly, years of education also showed moderate effects, with higher education levels associated with more stable or improved timing (e.g., Syllable duration variability), and voice-stability (e.g., Standard deviation shimmer), metrics.

Taken together, these findings show that although demographic, clinical, and cognitive covariates influence the magnitude and profile of speech changes, they do not account for the key intervention effects. The strongest group-level results—particularly Granularity in Animal Fluency and Main Tone in Routine, along with OSV and several voice-stability metrics—remained significant after adjustment, and Main Tone (Routine) retained its FDR-significant group effect (Nordic Walking vs Dance North). By contrast, more marginal results such as Average HNR (composite Dance South + Dance North vs Nordic Walking) and trend-level contrasts did not survive covariate adjustment. Thus, the core pattern of enhanced semantic precision and prosodic/voice-quality changes following dance remains statistically robust beyond demographic, cognitive, or disease-related differences.

## Discussion

This study aimed to assess the impact of an Amazonian Dance protocol, a culturally adapted intervention, on speech and language metrics in Brazilian PwP from two different Brazilian regions (North and South), compared to a time- and dose-matched Nordic Walking intervention. Dance groups showed significant improvements, especially in prosodic (Main Tone metric) and semantic organization (Granularity metric) features, particularly in the North region, where cultural familiarity with Amazonian dances was stronger. In contrast, participants in the Nordic Walking group showed declines in speech fluency and voice quality (Average HNR, Granularity and Speech Rate metrics). Even after covariate analysis the significant effect of dance interventions on prosodic and semantic features remained robust. These results provide novel evidence that dance interventions, particularly those culturally adapted, may modulate speech and language metrics in PwP.

Although dance interventions are well established for motor and psychosocial benefits in PD^44–46^ direct evidence for speech metrics has been scarce. The most directly comparable study is Karimi et al. (2025)^47^, who reported improvement in pitch stability in PwP after five years of weekly dance classes—consistent with our prosody improvements in the 12-week protocol. Other studies summarized in Theofanopoulou (2025)^34^ have documented gains in verbal fluency, lexical retrieval, and semantic clustering after dance training, though none clarified whether these reflected speech-motor or language-semantic processes. Our study results complement and extend this evidence showing improvements in both: prosodic control (Main Tone) and semantic precision (Granularity), quantified using automated speech and language analysis. This dual improvement suggests that dance can influence multiple layers of communication—from motor execution to lexical-semantic organization.

The more pronounced gains in the Amazonian Dance North group highlight the importance of cultural embodiment and regional identity. Participants’ familiarity with *Lundu* and *Carimbó*, traditional Northern Brazilian dances, may have amplified emotional engagement and motivation, fostering a stronger sense of belonging and expressive agency. Cultural embodiment may enhance adherence, self-expression, and social reward responses, all of which may facilitate deeper involvement in rhythmic, socially rich activities^34,48^. This finding aligns with recent meta-analyses emphasizing that culturally adapted and group-based performing arts interventions improve affective engagement and adherence^49–50^. In this context, Amazonian Dance may act as both a motor training and a sociocultural rehabilitation practice, engaging identity and emotional resonance as mediators of communicative vitality.

The observed improvements are also consistent with emerging mechanistic hypotheses regarding how dance influences speech-related brain pathways. Theofanopoulou (2025)^34^ proposed that dance training may affect not only limb-motor circuits but also the neural pathways controlling laryngeal and orofacial movements, supported by evidence that single neurons in the primary motor cortex can control both limb and vocal-motor actions. Neuroimaging studies across healthy adults and individuals with PD or MCI have shown dance-related changes in basal ganglia-premotor connectivity, frontal and temporal cortical plasticity, precentral gyrus structure, and networks spanning auditory, limbic, and sensorimotor areas^51–57^. Such findings suggest that dance can modulate distributed neural circuits relevant to not only speech-motor (e.g., prosody, timing), but also to non-motor/language (e.g., semantic organization) production.

The Nordic Walking group, by contrast, demonstrated declines in Granularity, HNR and Speech Rate, which may reflect natural progression of speech impairments in PD over the 12-week period including reduced voice quality, altered rhythm, and prosodic flattening^58–59^. These declines likely indicate the limited transfer of general aerobic exercise to speech-motor or language-semantic domains: although Nordic Walking is effective for improving gait and balance, it does not incorporate expressive and rhythmic components^44,60^, or auditory to motor integration, that possibly stimulate speech and language-relevant circuits. Thus, the contrasting patterns between Nordic Walking and dance strengthen the notion that dance interventions tap into neural systems beyond those targeted by conventional physical activity, yielding more direct benefits for communicative function.

Several covariates influenced the magnitude and profile of speech changes—particularly ethnicity, H&Y stage, sex, dance experience, cognition, and education—but none accounted for the core intervention effects. Ethnicity shaped variability and timing metrics; disease severity modulated whether participants improved more on timing versus voice-quality features; and female participants and those with prior dance experience tended to show stronger positive change in stability-related measures. Higher education and cognition modestly supported improvements in timing regularity. These patterns indicate that while individual characteristics shape responsiveness^61–62^, the dance-related gains in granularity and main tone remained robust and reproducible across analytic approaches. They reinforce that culturally grounded dance benefits extend beyond what can be predicted by demographic or clinical factors alone.

In conclusion, this study provides the first controlled evidence that a culturally adapted Amazonian Dance protocol can meaningfully modulate speech-motor and linguistic-semantics metrics in PwP. By combining automated speech and language analysis with a standardized 12-week intervention and a physically active control group, this work offers a reproducible framework for quantifying communication outcomes in dance research. The results highlight that culturally resonant, rhythm-based movement can enhance motor and non-motor aspects of speech/language—a promising direction for integrative, culturally sensitive rehabilitation in Parkinson’s disease.

## Study limitations

One limitation of this study is its pre–post intervention design without follow-up, which restricts the ability to draw conclusions regarding long-term retention or delayed intervention effects. Future trials should incorporate longitudinal follow-up assessments, including perceptual and patient-reported communication outcomes, to validate and expand upon these preliminary findings. Another limitation concerns the relatively narrow set of speech tasks and acoustic features employed. This restricted battery may not fully capture the complexity and heterogeneity of speech and language profiles in PwP. Future research should incorporate a broader range of ecologically valid tasks and more comprehensive linguistic-acoustic measures to enhance the characterization of dance-related changes in speech and language.

## Supporting information

TELL Codebook

## Resource availability Materials Availability

This study did not generate new unique reagents or materials.

## Data and Code Availability

Coded raw numerical data corresponding to TELL features and analysis data have been shared in Supplementary Tables. Fully raw data (i.e., voice recordings) cannot be shared because voice is a personally identifying information.

No custom code or software was generated or used in this study.

Any additional information required to reanalyze the data reported in this paper is available from the Lead Contact upon request.

## Acknowledgements

The authors acknowledge the participants in the study for their invaluable collaboration. Funding: This study was financed in part by the Federal University of Rio Grande do Sul (UFRGS) and Coordenação de Aperfeiçoamento de Pessoal de Nível Superior – Brasil (CAPES) – Finance Code 001. Constantina Theofanopoulou acknowledges funds from The Rockefeller University and New York University.

## Author contributions

Conceptualization: RAP, CT, ANH. Data Curation: RAP, CT, CCEGJ, MVA, PP. Formal Analysis: CT, RAP, MVA, AMS, PP. Funding Acquisition: ANH, CT. Supervision: ANH, CT. Writing – Original Draft Preparation: RAP, CT, ANH. Writing – Review & Editing: RAP, CCEGJ, MVAD, AMS, FGM, EPM, AG, ANH, CT. All authors read and approved the final manuscript.

## STAR METHODS

### Key Resources Table

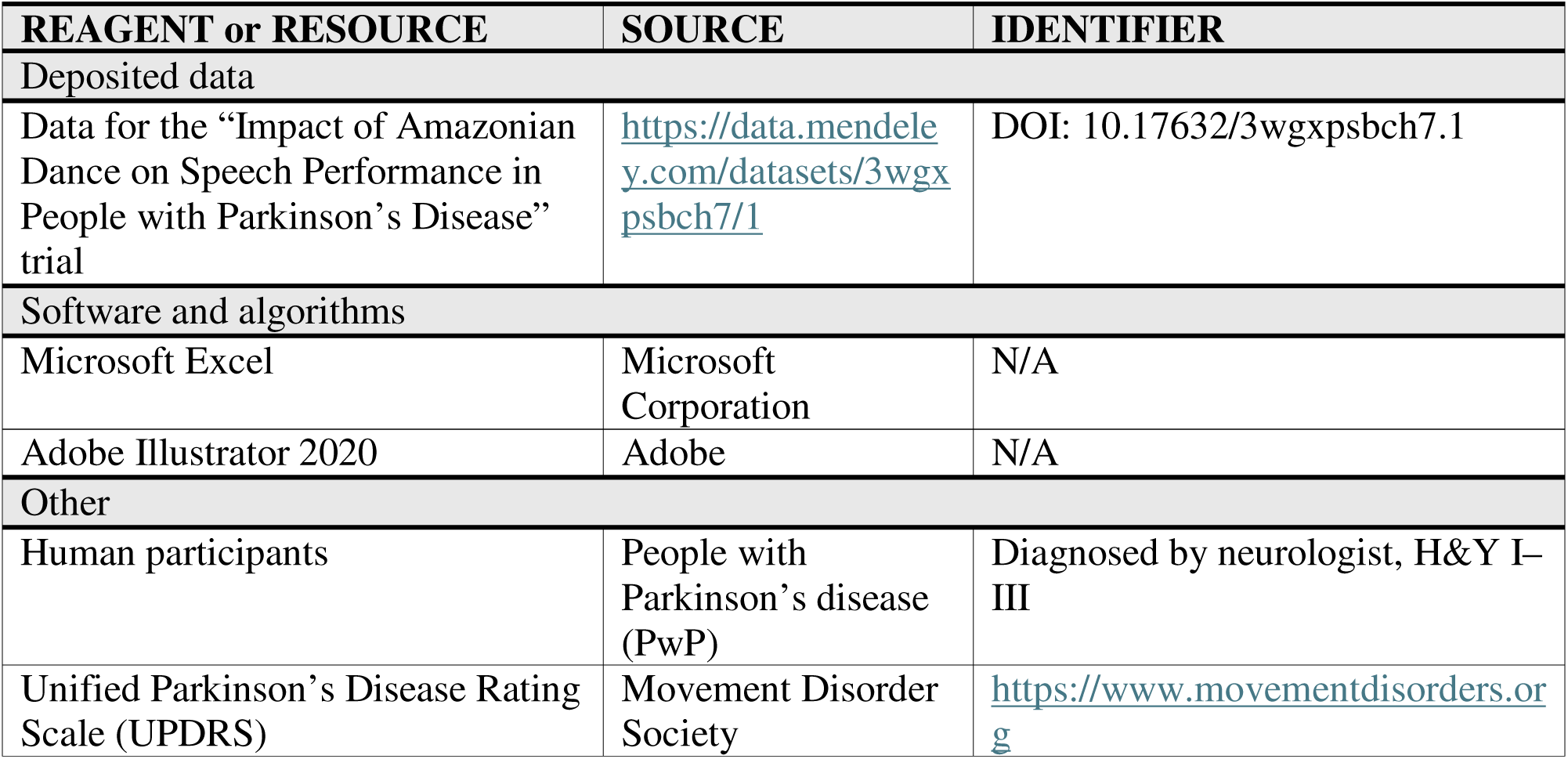

### Experimental model and study participant details

This study followed a randomized, parallel-group design approved by the Hospital de Clínicas de Porto Alegre (HCPA) Research Ethics Committee (CAAE: 75675023.9.0000.5327) and by the Federal University of Pará Ethics Committee (CAAE: 75675023.9.3002.0018). All the participants provided written informed consent prior to enrollment.

Sample size was determined following the protocol of a previously published study^24^ using GPower 3.1 software, based on an Analysis of Variance (ANOVA) - within/between interaction model (effect size = 0.40, α = 0.05, power = 0.95). The analysis indicated that a minimum of 16 individuals per group would be required. Allowing for an estimated 20% attrition rate, we targeted for enrollment of 20 individuals per group.

Participants were recruited from Dance and Nordic Walking Community Projects for PwP, Parkinson’s Associations, via social media and flyers distributed in two different regions of Brazil (North and South). Inclusion criteria were: a diagnosis of idiopathic PD, based on standard clinical diagnostic criteria determined by a neurologist^63^; age 50-80 years; both sexes; social-economically and racially diverse backgrounds; Hoehn and Yahr (H&Y) stages I-III; ≥ 1 year of stable medical treatment for PD with regular use of anti-parkinsonian drugs; ability to understand the verbal instructions for the tests and to walk with no walking aid; and native proficiency in Brazilian Portuguese with normal or corrected-to-normal vision and hearing. Exclusion criteria included the presence of other associated neurological or chronic diseases, recent surgery, or signs of Parkinson-plus syndromes.

The participants were allocated into three groups. Two groups received the Amazonian Dance intervention, delivered over 12 weeks, with two sessions per week, each lasting one hour. Dance-South Group (DS) comprised participants from Southern Brazil, where identification with Amazonian Dance is less prevalent, whereas Dance North Group (DN) comprised participants from Northern Brazil, who strongly identify with these regional cultural practices. The active control group, Nordic Walking Group (NW) completed a 12-week Nordic Walking intervention, also twice a week, with two one-hour sessions per week.

## Method details

### Amazonian Dance and Nordic Walking protocol

The instructor-led Amazonian dance and Nordic walking protocol were specifically designed for PwP, carried out in accessible and safe spaces and led by qualified instructors experienced in Parkinson’s population. The Amazonian Dance protocol was standardized and applied identically in both the South and North regions of Brazil. Nordic Walking was chosen as an active control condition because it is a well-established, low-cost, safe, and effective rehabilitation strategy for PwP^64^. The dose of both interventions—12 weeks, twice a week, 2 hours per week—was determined based on previous studies demonstrating optimal engagement and feasibility for older adults and PwP^65–66^. Detailed descriptions of the Amazonian Dance protocol^43^ and the Nordic Walking protocol^67^ have been published elsewhere.

**Figure 4.**
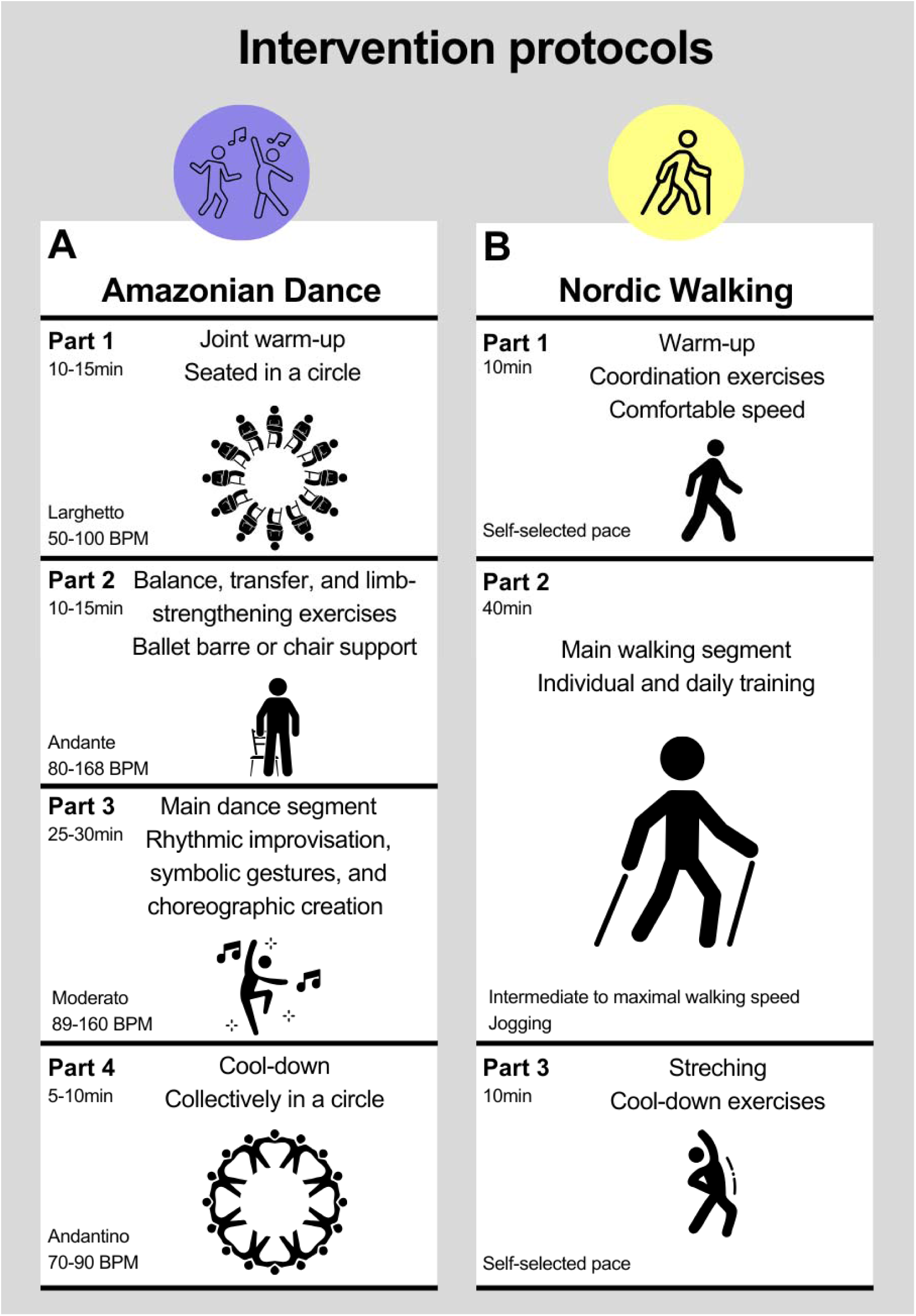
Structure of the Amazonian Dance and Nordic Walking intervention protocols. a, *Amazonian Dance protocol.* Each 60-minute session consisted of four sequential parts: Part 1 (10–15 min)—joint warm-up performed seated in a circle; Part 2 (10–15 min)—balance, transfer, and limb-strengthening exercises performed with support from a ballet barre or chair; Part 3 (25–30 min)—main dance segment incorporating rhythmic improvisation, symbolic gestures, and choreographic creation; and Part 4 (5–10 min)—cool-down performed collectively in a circle. Musical tempo varied across parts, progressing from *Larghetto* (50–100 BPM) to *Andante* (80–168 BPM), *Moderato* (89–160 BPM), and *Andantino* (70–90 BPM). The protocol was culturally adapted and standardized for both the Northern and Southern Brazilian sites. **b,** *Nordic Walking protocol.* Each 60-minute session consisted of three parts: Part 1 (10 min)—warm-up and coordination exercises at a self-selected pace; Part 2 (40 min)—main walking segment performed at intermediate to maximal walking speed or light jogging as tolerated; and Part 3 (10 min)—stretching and cool-down exercises at a self-selected pace.

### Instruments and Data Collection Procedures

All participants attended two data-collection sessions: a baseline evaluation (pre-intervention) and a post-intervention evaluation, after the 12-week program. To minimize motor-state fluctuations, all assessments were conducted during the medication "ON" state, between one to three hours after the participant’s regular dose of anti-parkinsonian medication. Pre- and post-intervention measurements were assessed at the same time of the day, while data collectors were blinded to the participants’ groups. Assessments were carried out by trained, experienced and qualified healthcare professionals.

Demographic and clinical data (age, sex, ethnicity, years of education, and years since the diagnosis) were collected using a structured anamnesis tool. The Brazilian version of the Goldsmith’s Dance Sophistication Index (Gold-DSI) was used to assess the participants’ dance experience^68^. The Gold-DSI comprises of 26 items assessing individual experiences of performing, watching, and knowing about dance. A Likert scale is used to classify the scores: not at all = 0; beginner = 1 to 2, intermediate = 3 to 4, advanced = 5 to 6, professional = 7. Global cognition was assessed using the Montreal Cognitive Assessment (MoCA), applying Brazilian education-adjusted cutoffs for mild cognitive impairment (MCI) and dementia^69^: level of education: ≥ 12 years – MCI (mild cognitive impairment) ≤ 22 score, dementia ≤ 21.5 score; 9-11 years - MCI ≤ 19.5 score, dementia ≤ 19.5 score; 5-8 years - MCI ≤ 19.5 score, dementia ≤ 16.5 score; 1-4 years - MCI ≤ 18 score, dementia ≤ 15.5 score; 0 years - ≤ 11.5 score, dementia ≤ 8.5 score. Disease stage was determined by the Hoehn and Yahr (H&Y) scale^70^, ranging from Stage I (unilateral involvement) to V (wheelchair-bound or bedridden), with higher stages indicating more advanced motor impairment.

### Speech data acquisition, processing and feature extraction

All speech and language data were collected and analyzed using the Toolkit to Examine Lifelike Language (TELL), a validated web-based platform that extracts acoustic, prosodic, and linguistic biomarkers sensitive to neurodegenerative disease. TELL provides automated, high-precision speech and language metrics based on naturalistic spoken responses^40–41^.

Participants completed two standardized speech tasks within the TELL app environment: 1) Animal Fluency: This is a widely used clinical measure of semantic memory and lexical retrieval. Participants were given 60 seconds to name as many different animal names as possible without repetition or derivational variants. The instructions for this task were as follows: ‘Name as many animals as you can in one minute. You may not repeat items, use proper names, or use words derived from the same root (e.g., table, tables, little table).” If the participant stopped speaking before the end of the minute, they were prompted with: “Can you think of more?”; 2) Routine Description: This task elicits spontaneous narrative speech for at least 60 seconds, requiring participants to describe their typical morning routine in detail. The instructions for this task were as follows: ‘Now tell me what a typical morning in your life is like. Give me as many details as you can. For example, instead of saying “I make breakfast” tell me everything you do to prepare breakfast. If the participant did not speak for at least one minute, we prompted them with: “Tell me more.”

All recordings were conducted individually with only the examiner and the examinee present, in a quiet room free of external noise. Participants were instructed to sit comfortably, silence electronic devices, remove jewelry that could interfere acoustically, and maintain a fixed distance from the microphone. An Apple iPad Air was used for speech recording, using its built-in microphone. Audio files were recorded and saved as single-channel Wave (.wav) files, with a sampling rate of 44.1 Hz and a bit-depth of 24 bits. Data collection was administered by trained healthcare professionals blinded to group assignment. Each recording underwent TELL’s five-step automated pipeline, encompassing volume normalization, voice activity detection, denoising, channel harmonization, and diarization^40–41^. Recordings were also transcribed via Whisper v.3.0 and manually revised by trained annotators following standardized criteria^40–41,71–72^.

The analyses focused on ten speech metrics, all shown to be sensitive to PD^16,72–73^: 1) Speech rate refers to the speed of speech production and is quantified in syllables per unit of time. Very high values may be associated with disinhibition, fluent aphasia, or manic episodes, whereas low values may indicate specific patterns of dysarthria or apraxia of speech, which are common in PD^74–76^; 2) Average pause duration captures the mean length of silences between sentences, words, or parts of words. Longer-than-normal pauses have been observed in conditions such as PD, AD, nonfluent primary progressive aphasia, depression, and schizophrenia, while sudden short pauses may be related to hypomanic states^77–79^; 3) Average syllable duration reflects the mean length of syllables in speech, with higher values suggesting motor speech difficulties commonly seen in PD, apraxia of speech, traumatic brain injury, and mild cognitive impairment^74,78,80–81^; 4) Pause duration variability captures fluctuations in the duration of pauses and may indicate impaired speech rhythm in conditions including PD, frontotemporal dementia, progressive nonfluent aphasia, amyotrophic lateral sclerosis, AD, depression, schizophrenia, and amnestic mild cognitive impairment^77–79^; 5) Syllable duration variability reflects variation in syllable length, and tends to be reduced in parkinsonian, ataxic, or right-hemisphere dysarthria, whereas increased variability has been reported in hypokinetic dysarthria^82–84^; 6) Harmonic-to-noise ratio (HNR) measures vocal efficiency by assessing the proportion of harmonic energy relative to noise, with low values suggesting asthenic voice or dysphonia, both being pathological voice conditions common in PD^85–86^; 7) Shimmer indexes cycle-to-cycle variability in amplitude, with increased values observed in PD and AD, reflecting irregular vocal fold vibration^87–88^; 8) Main tone refers to the predominant vocal pitch across the sample, with lower values often associated with depressive and neurodegenerative conditions^75,89–90^; 9) Semantic granularity reflects the degree of semantic specificity and content richness of the words produced, , ranging from broad and unspecific terms (e.g., “entity,” “thing”) to more fine-grained categories (e.g., “animal,” “dog,” “bulldog”). Low values may indicate impaired access to semantic memory^73,91–92^; 10) Ongoing semantic variability (OSV) quantifies the semantic distance between consecutive words and examines the variability of these distances across the narrative. High variability suggests flexible semantic exploration, whereas low variability indicates restricted or stereotyped semantic production^91–92^. Together, the first eight features primarily index speech-motor and phonatory control, capturing articulatory timing, prosodic rhythm, and vocal stability; in contrast, the final two features—semantic granularity and ongoing semantic variability—reflect higher-order language and semantic processing, providing complementary information about lexical access, semantic organization, and discourse structure.

### Quantification and statistical analysis

Descriptive statistics were first computed for all demographic and clinical variables and are reported as means, standard deviations and proportions. Pre- and post-intervention data from two speech tasks, Animal Fluency and Routine, were analyzed to examine changes in speech performance across three groups: Dance South (DS), Dance North (DN) and Nordic walking (NW). Each participant contributed data at two timepoints (pre- and post-intervention), and for each speech metric individual-level change score (Δ = post − pre) was computed. Positive Δ values indicated increases and negative values indicated decreases in the measured metric. Importantly, whether a positive Δ reflected functional improvement depended on the nature of the metric: for example, higher speech rate or higher Harmonics-to-Noise Ratio (HNR) are considered improvements (so positive Δ is beneficial), whereas lower average pause duration and lower shimmer variability are considered improvements (so negative Δ is beneficial). These directional expectations were established a priori based on the known psychometric properties of each feature and are summarized in the TELL Speech Feature Codebook (**Document S1**).

To run analysis at multiple levels, the data was first organized in a long-format within-subject table containing, for each participant, task, and metric, their pre, post, and Δ values (**Table S1**). Because each metric was collected separately during the Animal Fluency and Routine tasks, in addition to individual task analyses, a cross-task composite value for each metric by averaging the scores from the two tasks for each participant was computed, producing an individual-level table of composite pre, post, and Δ values (**Table S3**). These individual-level tables served as the basis for all subsequent group-level analyses.

Within each group, pre-post changes were tested using paired-samples t-tests when normality assumptions were met, and Wilcoxon signed-rank tests otherwise. These analyses were performed both at the task level (separately for Animal Fluency and Routine) (**Table S2**) and at the composite level (combining the two tasks) (**Table S4**). For each group × task × metric combination, the number of paired observations were reported, pre and post means with 95% confidence intervals (CIs), the mean Δ and its 95% CI, the test statistic, the p-value, and the effect size as Cohen’s d (for t-tests).

To compare the magnitude of change across groups, between-group analyses were conducted on the individual Δ scores. These tests assessed all pairwise group contrasts (DS vs NW, NW vs DN, DS vs DN) as well as a combined contrast comparing DN+DS (the two dancing groups) versus NW (the walking group). Analyses were performed separately for the two tasks (**Table S5**) and for the composite scores (**Table S6**). Welch’s t-tests were applied when normality assumptions were met, and Mann-Whitney U tests otherwise. For each contrast, the group means of Δ with 95% CIs, the test statistic, uncorrected p-value, and effect sizes as Hedges’ g (for mean differences) and rank-biserial r (for nonparametric contrasts) were reported. All within- and between-group p-values were corrected for multiple comparisons using the Benjamini-Hochberg false discovery rate (FDR) procedure, and the FDR-adjusted q-values are reported alongside the uncorrected p-values.

To examine potential sources of inter-individual variability in treatment response, demographic, cognitive, and clinical covariates were further tested for associations with speech changes. The covariates analyzed were Sex, Ethnicity, Dance Experience, Global Cognition, Years of Education, Age, and disease stage. Multivariable general linear models (ANCOVA-style) were first fitted to predict Δ scores from Group and all seven covariates entered simultaneously. For each factor, the F-statistic, uncorrected p-value, FDR-corrected q-value, and partial eta-squared (η²), as well as the model R² (**Tables S7**) were reported. To isolate the contribution of each covariate without adjustment for the others, a series of single-predictor models including Group and only one covariate at a time, reporting the F-statistic, p-value, and partial η² for the covariate (**Table S8**), were also ran.

Finally, the descriptive pattern of speech changes across levels of categorical covariates and as a function of continuous covariates were summarized. For the categorical covariates (Sex, Ethnicity), the mean pre, post, and Δ scores within each subgroup (**Table S9**) were computed. For the continuous covariates (Dance Experience, MoCA, Years of Education, Age, H&Y), simple linear regressions of Δ were fitted on each covariate separately, and the regression slope (β), Pearson correlation coefficient (r), and p-value were reported (**Table S10**). All analyses were conducted using two-sided tests. Confidence intervals reflect 95% coverage.

## Adidtional resources

This study was conducted according to CONSORT 2025 Statement: updated guideline for reporting randomized trial^93^ (Supplementary material – CONSORT checklist).

## Supplemental information index

**Document S1.** Tell Codebook

**Tables S1-S10.** Excel including Tables S1-S10

**Table S1 -** Individual pre–post data per participant, task, and speech metric.

**Table S2 -** Within-group pre–post analyses of speech metrics for Animal Fluency and Routine tasks.

**Table S3 -** Individual composite pre–post data per participant and speech metric.

**Table S4 -** Within-group pre–post analyses of composite speech metrics.

**Table S5 -** Between-group comparisons of pre–post change (Δ) on speech metrics.

**Table S6 -**Between-group comparisons of pre–post change (Δ) on composite speech metrics.

**Table S7 -** Single-predictor models testing the effect of individual covariates on pre–post change (Δ).

**Table S8 -** Multivariable models testing Group and covariate effects on pre–post change (Δ).

**Table S9 -** Mean pre, post, and change (Δ) scores by categorical covariate levels.

**Table S10** - Slopes of pre–post change (Δ) regressed on continuous covariates.

**CONSORT checklist**

