## Supplementary material for "Impact of Amazonian Dance on Speech Performance in People with Parkinson’s Disease": TELL Codebook

**The source of this information is the explanation provided in TELL app (https://tellapp.org/) regarding each metric, the authors just organized it in one unique file.**

- **Speech Rate**

Brief explanation:

This metric indicates the speed of speech production, quantified in syllables per unit of time. Very high values may be associated with disinhibition, fluent aphasia or manic episodes. Low values may indicate specific patterns of dysarthria or apraxia of speech.

Full explanation:

Speech rate derives from the number of syllables produced per unit of time. The higher the value, the faster the speech. This metric is one of the best predictors of subjective fluency in research on speech and language disorders (Cannizzaro et al., 2004; Covington, 2005; Cucchiarini et al., 2002; Feyereisen, Pillon, & De Partz, 1991; Kormos & Dénes, 2004; Shenker, 2006) and is sensitive to multiple underlying alterations that can be motor, cognitive, or affective. For example, speech rate decreases in cases of progressive non-fluent aphasia, Parkinson's disease, Alzheimer's disease (Ash et al., 2007; Carlomagno et al., 2005; Martínez-Sánchez et al., 2013; Meilán et al., 2014; Sajadi et al., 2012) and mild cognitive impairment (Gosztolya et al., 2018), but may increase in psychotic states due to schizophrenia, anxiety or manic states of bipolarity.

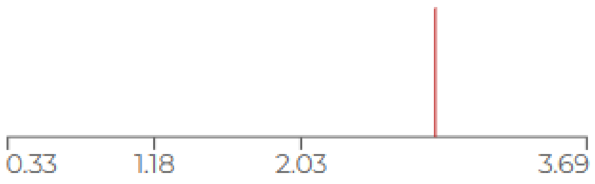


- **Average pause duration**

Brief explanation:

This metric captures the length of silences between sentences, words, or parts of words. Longer-than-normal silences are observed in a number of conditions, including Parkinson's disease, Alzheimer's disease, nonfluent primary progressive aphasia, depression, and schizophrenia. The occurrence of sudden shorter silences may be associated with hypomanic episodes.

Full explanation:

The duration of silences is influenced by various mechanisms. The appearance of longer silences may be due to motor difficulties during speech, cognitive deficits in finding the desired words, or affective changes. This metric is a robust marker of several neurological and psychiatric conditions, including Alzheimer's disease (Singh et al., 2001), non-fluent primary progressive aphasia (García et al., 2022), depression, and schizophrenia (Rapcan et al., 2022). Its accurate interpretation requires additional information from neurological, neuropsychological, or neuropsychiatric scales.

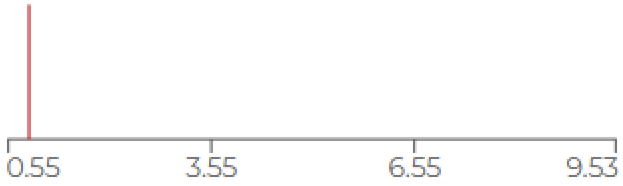


- **Average syllable duration**

Brief explanation:

This metric indicates the average length of syllables in speech. Higher values may reflect motor difficulties in speech in patients with Parkinson's, apraxia, traumatic brain injury, and mild cognitive impairment, among others.

Full explanation:

The syllable is a basic representative unit of the rhythmic organization of speech (Ziegler et al., 1993). It is measured as the phonation time (phonation time) divided by the number of syllables identified in speech. A shorter syllable duration is an indicator of normal pronunciation for spontaneous speech (Table et al., 2021). Syllable duration depends on the number of phonemes and the emphasis given to them (Crystal & House, 1990), but can also be affected by pathologies that hinder speech production, including Parkinson's

(Martínez-Sánchez et al., 2014), apraxia (García et al., 2022), traumatic brain injury (Campbell et al., 1995), and mild cognitive impairment (Themistocleous, 2020).

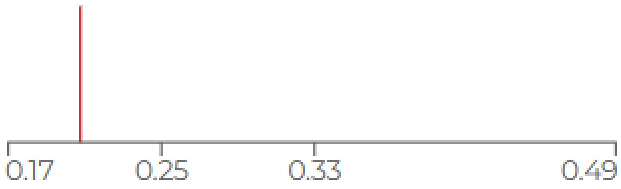


- **Pause duration variability**

<u>Brief explanation:</u>

An increase in the variability of pause duration may be indicative of conditions characterized by disorders affecting speech rhythm, including frontotemporal dementia, progressive nonfluent aphasia, amyotrophic lateral sclerosis, Alzheimer's, and amnestic mild cognitive impairment.

<u>Full explanation:</u>

The length of silences is influenced by various mechanisms. The appearance of longer silences may be due to motor difficulties during speech, cognitive deficits in finding the desired words, or affective changes. This metric is a robust marker of several neurological and psychiatric conditions, including Alzheimer's disease (Singh et al., 2001), primary progressive nonfluent aphasia (García et al., 2022), depression, and schizophrenia (Rapcan et al., 2010, 2010). Its accurate interpretation requires additional information from neurological, neuropsychological, or neuropsychiatric scales.

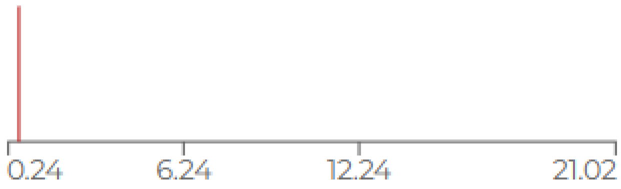


- **Syllable duration variability**

<u>Brief explanation:</u>

Variation in syllable duration is reduced in cases of ataxic, parkinsonian, or right hemisphere dysarthria (Kent & Rosenbeck, 1982; Ludlow et al., 1987). In other cases, such as hypokinetic dysarthria, its value has been observed to increase (Brenk, 2018).

Full explanation:

The syllable is a basic representative unit of the rhythmic organization of speech (Ziegler et al., 1993). It is measured as the phonation time (phonation time) divided by the number of syllables identified in speech. A shorter syllable duration is an indicator of normal pronunciation for spontaneous speech (Table et al., 2021). Syllable duration depends on the number of phonemes and the emphasis given to them (Crystal & House, 1990), but can also be affected by pathologies that hinder speech production, including Parkinson's (Martínez-Sánchez et al., 2014), apraxia (García et al., 2022), traumatic brain injury (Campbell et al., 1995), and mild cognitive impairment (Themistocleous, 2020).

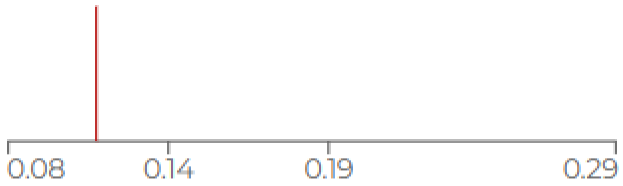


- **Granularity**

Brief explanation:

Granularity is a measure of how specific the words used by the participant are. It ranges from "entity" or "thing" as the lowest granularity words, to progressively more specific words (e.g., "animal", "dog", "bulldog"). A low granularity value may indicate loss of cognitive deficits.

Full explanation:

Semantic granularity is obtained through a hierarchical tree of nodes that progresses from the highest hypernym ('entity') to progressively more specific concepts (e.g., 'animal', 'dog', 'bulldog'). It is defined as the number of nodes between a word and its related 'entity' (e.g., words with low granularity are closer to the 'entity' than words with higher granularity), indicating that the former represents more general concepts. People with Alzheimer's disease show a reduction in semantic granularity, favoring conceptually broad words over specific ones (e.g., 'flower' over 'rose') (Ferrante & Migoeot et al., 2023; Sanz et al., 2022). This pattern may reflect memory and/or executive dysfunctions (e.g., a reduced capacity for abstraction), as well as neurofunctional disruptions in temporal regions (Chhatwal et al., 2018) associated with conceptual accuracy (Beaty et al., 2018).

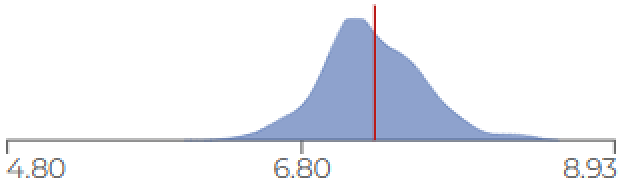


- **OSV (Ongoing semantic variability)**

<u>Brief explanation:</u>

OSV calculates the semantic distance between pairs of consecutive words and considers the overall variability of this distance in the text. On the one hand, a very high value may reflect a lack of coherence in the discourse, and on the other hand, a low value may reflect deficits in the suppression of previous semantic information, possibly due to inhibitory alterations.

<u>Full explanation:</u>

Semantic variability refers to changes in conceptual distance between successive words. Compared to healthy individuals, people with Alzheimer's mention semantically more distant words (Sanz, C. et al., 2022), unlike people with Parkinson's (Arias-Trejo, N. et al., 2021), who also show an increase in the semantic priming effect (Marí-Befffa, P., 2005) and less semantically different groups (Raskin, S.A., et al., 1992). Because semantic variability requires suppressing the current semantic field to activate another, these patterns may reflect weak semantic inhibition, a typical feature of Parkinson's (Siquier, A., et al., 2021; Castner, J.E. et al., 2007).

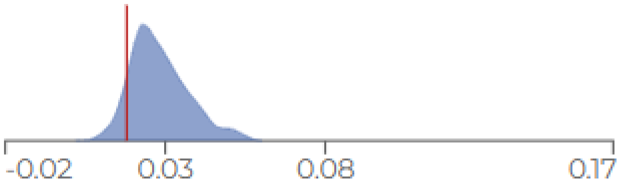

- **Averege HNR**

Brief explanation:

The harmonic-to-noise ratio (HNR) is a measure of speech efficiency, as it reflects the ability to transform air from the lungs into vibration of the vocal cords. Low values may be indicators of pathologies, including asthenic voice and/or dysphonia.

Full explanation:

The harmonic-to-noise ratio (HNR) is a measure of voice quality calculated by dividing the energy of voice harmonics by the energy of background noise. It is expressed in dB and is a measure of speech efficiency—the ability to transform air from the lungs into vocal cord vibrations. Low HNR values are indicators of asthenic voice and/or dysphonia (Teixeira et al., 2013). This parameter has been used to identify movement disorders through speech analysis (Dao et al., 2022). HNR can also be used as a marker to detect differences between healthy and fatigued voices due to sleep deprivation (Kumar et al., 2023).

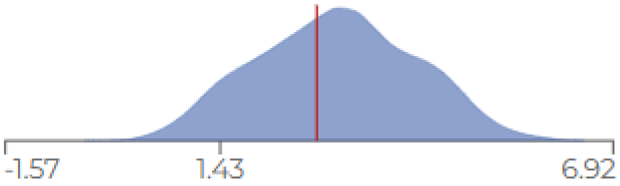


- **Standard deviation HNR**

Brief explanation:

Greater variability in this metric is associated with conditions that include dysphonia, vocal cord paralysis and chronic laryngitis.

Full explanation:

The harmonic-to-noise ratio (HNR) is a measure of voice quality calculated by dividing the energy of voice harmonics by the energy of background noise. It is expressed in dB and is a measure of speech efficiency—the ability to transform air from the lungs into vocal cord vibrations. Low HNR values are indicators of asthenic voice and/or dysphonia (Teixeira et al., 2013). This parameter has been used to identify movement disorders through speech analysis (Dao et al., 2022). HNR can also be used as a marker to detect differences between healthy and fatigued voices due to sleep deprivation (Kumar et al., 2023).

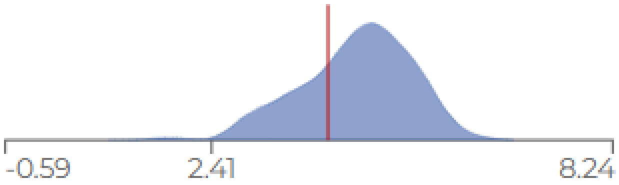


- **Shimmer**

Brief explanation:

Shimmer is the variation in amplitude from peak to peak (local maxima). Different parameters derived from this descriptor are used to measure the impact of Parkinson's disease on the human voice, and they have been found to be greater in people with Alzheimer's disease.

Full explanation:

Shimmer is impacted by changes in glottis resistance and vocal cord injuries and is related to the presence of noise and broken breathing (Teixeira et al., 2013). Different parameters derived from this descriptor are used to measure the impact of Parkinson's disease on the human voice (Azadi et al., 2021), and they have been found to be higher in people with Alzheimer's disease (Hason & Krishnan, 2022).

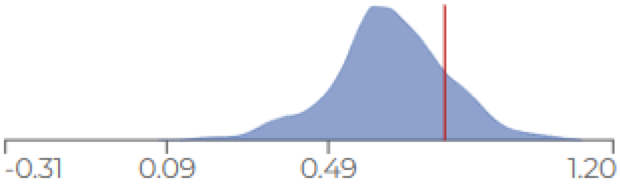


- **Standard deviation shimmer**

Brief explanation:

Its variation is associated with problems in control over the vocal tract and injuries to the vocal cords.

Full explanation:

Shimmer is impacted by changes in glottis resistance and vocal cord injuries and is related to the presence of noise and broken breathing (Teixeira et al., 2013). Different parameters derived from this descriptor are used to measure the impact of Parkinson's disease on the human voice (Azadi et al., 2021), and they have been found to be higher in people with Alzheimer's disease (Hason & Krishnan, 2022).

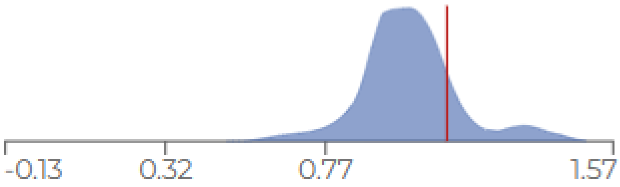


- **Main tone**

Brief explanation:

Low values of the main tone are associated with features present in depressive or neurodegenerative conditions, including the behavioral variant of frontotemporal dementia and the nonfluent variant of primary progressive aphasia.

Full explanation:

In certain neuropsychiatric conditions, the fundamental tone of speech (pitch) is associated with depressive traits: the lower the fundamental tone values, the greater the expression of depressive symptoms. This metric has also been useful in predicting the occurrence of initial psychotic episodes. Other studies have documented a decrease in the fundamental tone of speech in neurodegenerative conditions such as behavioral variant frontotemporal dementia and nonfluent variant primary progressive aphasia.

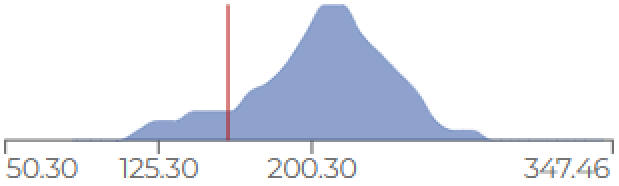


**Support articles:**

Toolkit to Examine Lifelike Language v.2.0: Optimizing Speech Biomarkers of Neurodegeneration

Toolkit to Examine Lifelike Language (TELL): An app to capture speech and language markers of neurodegeneration
